# Neuron-derived Thioredoxin-80: a novel regulator of type-I interferon response in microglia

**DOI:** 10.1101/2022.03.09.483648

**Authors:** Julen Goikolea, Jean-Pierre Roussarie, Gorka Gerenu, Raul Loera-Valencia, Maria Latorre-Leal, Angel Cedazo-Minguez, Patricia Rodriguez-Rodriguez, Silvia Maioli

**Author notes:** Corresponding author: Julen Goikolea. These authors contributed equally to this work (Patricia Rodriguez-Rodriguez, Silvia Maioli).

## Abstract

Oxidative stress and neuroinflammation play a central role in Alzheimer’s Disease (AD) pathogenesis. However, the mechanism by which these processes lead to neurodegeneration is still not fully understood. Thioredoxin-1 (Trx1) is an antioxidant protein that can be cleaved into a peptide known as Thioredoxin-80 (Trx80), which modulates monocyte function in the periphery and shows anti-amyloidogenic properties in the brain. In this study we aimed to further clarify the biological function of this peptide and its regulation in the brain. We show that neurons are the main producers of Trx80 in the brain. Trx80 levels increase *in vivo* both in normal aging and in young APP^*NL-G-F*^ mouse model of amyloid pathology. Trx80 levels were increased in neurons in primary culture treated with either rotenone or 27-hydroxycholesterol, what suggests that Trx80 production is stimulated upon oxidative stress. RNA-sequencing followed by differential gene expression analysis revealed that Trx80 induces microglia activation into a phenotype compatible with interferon response microglia. Finally, we determine that the induction of this microglia phenotype by Trx80 is Trem2-dependent. This study identifies Trx80 as a novel neuron-derived signaling mechanism that modulates microglia function under stress conditions. Strategies to regulate Trx80 levels could be beneficial against AD pathology.

## Introduction

Microglia are the resident immune cells of the central nervous system and play important roles in brain development and physiology (1). Their relevance in disease pathogenesis has gained increasing interest in AD, where genome-wide association studies (GWAS) have established that many AD genetic risk variants are confined to microglia transcriptional enhancer regions (2). Moreover, rare genetic variants encoding triggering receptor expressed on myeloid cells 2 (TREM2), which plays an essential role as a regulator of microglia function, have been identified as significant genetic risks factors for late onset AD (3-5). During AD pathology, a diversity of microglial cell states has been identified (6-8). Of special interest are two different, mutually exclusive, reactive microglia states present in normal aging that are modified by AD risk factors *in vivo*: Activated Response Microglia (ARM) and Interferon Response Microglia (IRM)(9). ARMs show overexpression of genes of the Major Histocompatibility complex type II (MHCII) and they are enriched in AD risk genes whereas IRMs are characterized by an upregulation of type-I interferon response genes. The mechanisms that govern the appearance of these different microglia states and their role in neurodegeneration remain to be fully understood and could provide new avenues for therapeutic intervention.

Thioredoxin-1 (Trx1) is a highly conserved endogenous dithiol with functions that span from maintenance of cellular redox homeostasis to chemokine activity (10, 11). Trx1 can be cleaved into an 80 amino acid long peptide, known as Thioredoxin-80 (Trx80) (12). Trx80 function has been widely investigated in the periphery, where it is secreted and triggers innate immunity and proinflammatory responses (13, 14).

Our group reported that Trx80 inhibits Aβ-induced toxic effects in cell cultures as well as Aβ42 accumulation in the brain (15). Moreover, Trx80 expression rescued the lifespan and locomotor impairments observed in Aβ42-expressing *Drosophila melanogaster* models (16). Interestingly, Trx80 protein levels are depleted in brains of AD patients (15). Nevertheless, the biological function and mechanisms that regulate Trx80 production and secretion in the brain are still unknown.

In the present study we show that Trx80 levels in the brain are affected by aging and amyloid pathology, two important drivers of AD pathology. Furthermore, we identify that oxidative stress, a converging event for aging and amyloidosis increases Trx80 production by neurons. Finally, we show that Trx80 leads to microglia differentiation into a type-I-interferon response transcriptional profile in a Trem-2 dependent manner. Together, our results unveil a new mechanism that regulates microglia function with implications for AD pathology.

## Materials and Methods

### Animals

Male C57BL/6J wild-type mice at 2, 8 and 22 months of age were obtained from Janvier (Janvier labs, Le Genest-Saint-Isle, France). Female 3 and 10 months old homozygous APP^NL-G-F^ knock in mice and their wild-type C57BL/6J (WT) controls were available at our department at KI. Male 22 months old Cyp27A1 overexpressing mice and their littermate WT controls were available in our group. Mice were kept under controlled temperature (21±1°C) and humidity (55±5%) on a 12-h light-dark cycle and food/water were provided ad libitum.

### Cortico-Hippocampal Mouse Primary Neurons and Microglia cultures

Cortico-Hippocampal primary neuronal cultures from E17 C57BL/6J mouse embryos (Janvier labs) were performed as previously reported (17). Briefly, dissected tissue was trypsinized in 0.25% trypsin, 1mM EDTA (T4049, Merck, Darmstadt, Germany) in EBSS (#14155063, Thermo Fisher Scientific, Waltham, Massachusetts, USA) for 15 minutes at 37°C. After centrifugation (500 x g, 5min) the tissue was mechanically disintegrated in a solution containing 170 ug/ml DNAse (#10104159001, Merck) in EBSS. The supernatant was transferred to a fresh tube, centrifuged (500 *x* g, 5 minutes) and the pellet was resuspended in serum-free neurobasal medium (#21103049, Thermo Fisher Scientific) with B-27 (#17504044, Gibco, Thermo Fisher Scientific), Penicillin/Streptomycin (#15140122, Thermo Fisher Scientific) and Glutamax (#35050061, Thermo Fisher Scientific). Cultures were incubated at 37°C and 5 % CO_2_.

Microglia and astrocyte cultures were performed following the procedure described for neurons, but cells were resuspended in DMEM/F-12, GlutaMAX (#10565018, Thermo Fisher Scientific) and supplemented with 10% Heat-inactivated FBS (#10500064, Gibco, Thermo Fisher Scientific) and Penicillin/Streptomycin. Cultures were incubated at 37°C and 5 % CO_2_ for 24h before complete media change. Further media changes were performed every 3 days until confluency was reached. Microglia was dissociated from mixed glial cultures by mild-trypsinization (0.25% trypsin, 1 mM EDTA diluted 1:4 in DMEM/F-12, GlutaMAX serum-free) as previously reported (18).

### Cell treatments

All neuron treatments were performed at 10 days in vitro (DIV). Neurons were treated 1 μM 27-hydroxycholesterol (27-OHC, Avanti Instruchemie, The Netherlands) or vehicle (DMSO, Sigma) for 2h before collection. 100 nM Rotenone (#R8875, Merck) or vehicle (DMSO) for 18h before collection. 1 μM Trx80 (#2329-TX, R&D Systems, Minneapolis, Minnesota, USA) or vehicle (PBS) for 24h before collection.

Microglia cultures were kept at 37°C and 5 % CO_2_ for 24h before treatment. Cells were treated with 1 μM Trx80 peptide or vehicle (PBS) for 24h before collection.

### Sample preparation and western blot

Immunoblotting was performed as previously described (17). Briefly, mouse brain tissue and cell lysates were homogenized in 50 mM Tris-HCL, 150 mM NaCl, 1% Triton-X containing phosphatase and protease inhibitors (Sigma-Aldrich, St. Louis, Missouri, USA). Homogenates were run in polyacrylamide gels (Bio-Rad, Hercules, California, USA and transferred to BioTraceTM Nitrocellulose membranes (GE healthcare, Chicago, Illinois, USA). Membranes were blocked for 1h in 5% skim milk in TBS-Tween-20 (TBS-T) prior to overnight incubation with primary antibody at 4°C. Primary antibodies: anti-Trx80 mouse (7D11 #11543, Cayman chemicals, Ann Arbor, Michigan, USA), anti-Trx1 goat (#11538 Cayman chemicals), anti-NRF2 rabbit (ab137550, Abcam, Cambridge, United Kingdom), Anti-β-Actin mouse (AC-15, Sigma-Aldrich), anti-GAPDH mouse (6C5, ab8245, Abcam) and Anti-α-Tubulin antibody (T9026, Sigma-Aldrich). Fluorescent secondary antibodies (LI-COR Biosciences, Lincoln, Nebraska, USA) were used for 2h at room temperature and visualized using ODYSSEY Infrared Imaging System (LI-COR Biosciences). Band intensity signal was quantified by ImageJ software. Each band signal value was normalized against loading controls (tubulin/actin/GAPDH signal value).

### RNA extraction and Real-time quantitative PCR

Total RNA extraction, reverse transcription, and real-time quantitative PCR (RT-qPCR) amplification was performed using the TaqMan Fast Advanced Cells-to-Ct Kit following the manufacturer’s instructions (Thermo Fisher). Briefly, cells were washed in PBS and lysed in solution containing DNase for 5 minutes at room temperature. Lysis was terminated at room temperature by a 2-minute incubation with Stop Solution. Lysates were reverse transcribed in a SimpliAmp Thermal cycler (Life Technologies, Carlsbad, California, USA). Quantification of selected genes was done using specific primers for *Ifit2, Ifit3, Irf7, Spp1, Axl, Trem2* and *Gapdh* (Life Technologies) in a 7500 Fast Real-Time PCR system (Applied Biosystem, Life Technologies). GAPDH was used as housekeeping gene.

### Small Interference RNA Transfection

*Trem2* and *Axl* knockdown were performed using siRNA designed by Horizon Discovery (Accell Mouse Trem2 siRNA SMARTpool, E-040918-00-0005; Accell Mouse Axl siRNA SMARTpool, E-040941-00-0005). Mouse primary microglia (80% confluence) were transfected with a final concentration of 0.5 μM per well for 8h according to the manufacturer’s instructions. The efficiency of the knockdown were measured using mRNA levels of *Trem2* and *Axl*.

### Immunofluorescence

Microglial cells were fixed in 4% PFA for 10 min followed by permeabilization and washing steps (0.25% Triton-X in PBS). Cells were then incubated in 5% BSA Blocking buffer 1 h at room temperature. As primary antibodies, anti-Ifit2 rabbit (PA3-845, Sigma-Aldrich), anti-ISG15 rabbit (PA5-79523, Thermo Fisher Scientific) anti-OPN goat (AF808-SP, R&D systems), anti-Iba-1 rabbit (#019-19741, Wako, Japan) and anti-Iba-1 goat (NB100-1028, NOVUS) were incubated overnight at 4°C. Thereafter, samples were washed three times and incubated 1h at room temperature with secondary antibodies: Alexa Fluor Plus 488 goat anti-Rabbit (A32731, Thermo Fisher Scientific) and Alexa Fluor Plus 546 donkey anti-goat (A11056, Thermo Fisher Scientific). Identification of the nuclei of cell bodies was performed using 4, 6 diamino-2-phenylindole (DAPI; Sigma).

Immunofluorescence of mice cortical samples was performed as previously described (19). The following primary antibodies were used: anti-Iba-1 goat and anti-Ifit2 rabbit. Sections were then incubated for 2 h at room temperature with the secondary antibodies Alexa Fluor Plus 488 goat anti-Rabbit and Alexa Fluor Plus 546 donkey anti-goat. Identification of the nuclei of cell bodies was performed using DRAQ5 fluorescent prove solution (#62251, Thermo Fisher Scientific). Omission of the primary antibody was done as a control staining. Sections were mounted using fluorescence mounting medium (DAKO Cytomation, Glostrup, Denmark). Images were obtained in a Zeiss LSM900-Airy confocal microscope performing a Z-stack at cortical areas. Fluorescence intensity of Isg15 per microglial cell was measured by the ImageJ 1.383 software (NIH, MA, USA).

### RNA sequencing

Library preparation and RNA sequencing was outsourced to the Bioinformatics and Expression Analysis (BEA) core facility at The Karolinska Institute. Briefly, total RNA was subjected to quality control with Agilent Bioanalizer 2100 using an RNA pico chip according to the manufacturer’s instructions. cDNA libraries were prepared using the Truseq stranded mRNA samples preparation protocol (Illumina, San Diego, California, USA), which includes mRNA isolation, cDNA synthesis, ligation of adapters and amplification of indexed libraries. The yield and quality of the amplified libraries was analyzed using Qubit (Thermo Fisher Scientific) and the Agilent TapeStation. Libraries were then normalized, multiplexed and sequenced on the Illumina Nextseq 550 for a 75-cycle v2 sequencing run generating 75 bp single-end reads.

### Statistical analysis

All experiments were performed at least in triplicate. Statistical analysis was performed using Prism software (GraphPad 9). Student’s t-test was used to evaluate the significance of the statistical difference between two groups. The statistical differences among three groups was calculated using One-way analysis of variance (ANOVA). Fisher’s exact test was used to assess the gene expression overlap between Trx80-activated microglia, IRMs and ARMs. For all tests, P < 0.05 was considered statistically significant.

## Results

### Trx80 is produced by neurons and its levels are increased with normal aging and in young Appki^*NL-G-F*^ mouse

The relative contribution of the different major cell types in the brain to the Trx80 pool is unknown. To clarify this issue, we determined Trx80 protein levels in mouse primary cultures of neurons, astrocytes, and microglia. Our results show that neurons display higher levels of Trx80 than astrocytes and microglia (p<0.005 and p<0.001, respectively) (**Figure 1a**). These results agree with previous reports that found most Trx80 immunoreactivity in pyramidal neurons in the human brain (15). Protein levels of its precursor Trx1, were, however, comparable between these cell types. RT-qPCR analysis revealed that *Txn1* (the gene that codes for Trx1) expression is higher in neurons than in astrocytes or microglia (p<0.0001) (**Figure 1b**). These results are in agreement with publicly available *in vivo* single-cell transcriptomics data (**Figure 1c**, Allen Institute for Brain Science (2020), celltypes.brain-map.org/rnaseq (20)). Together, these data suggests that, as opposed to the other cell types, a big proportion of Trx1 in neurons is transformed into Trx80.

**Figure 1.**
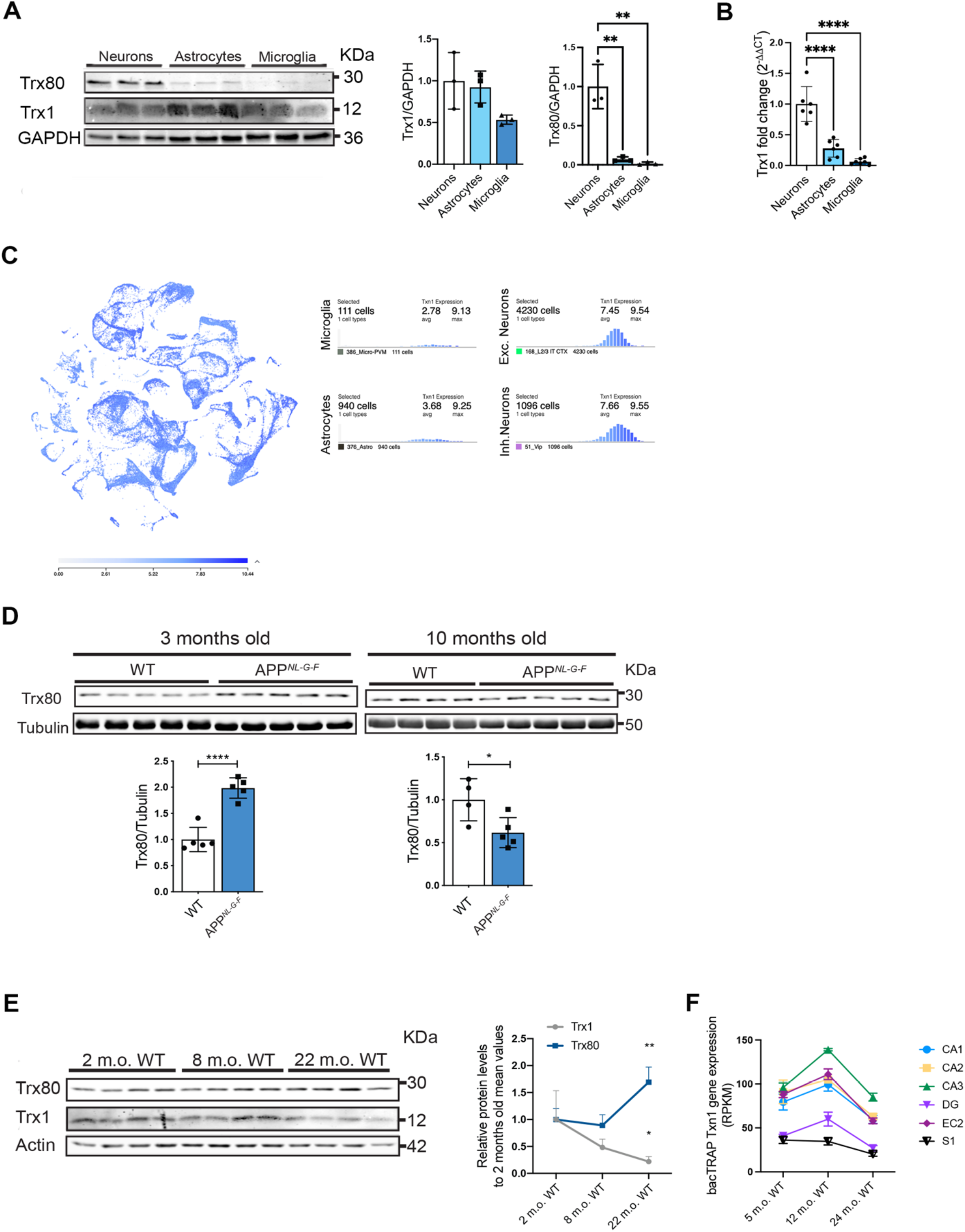
Trx80 is produced by neurons and its levels are increased with normal ageing and in young Appki^*NL-G-F*^ mouse model of amyloid pathology. **A)** Western-blot analysis of mouse primary neurons, astrocytes and microglia. Trx80, Trx1, GFAP and Iba-1 band intensities are normalized against GAPDH (n=3 biological replicates). Results are expressed as mean ± SD. **B)** RT-qPCR analysis of Txn1 gene expression levels in mouse primary neurons, astrocytes and microglia. Values were normalized against each samples *Gapdh* gene expression levels. **C)** *Txn1* gene expression from *in vivo* single-cell transcriptomics data Allen brain map transcriptomics explorer. **D)** Western-blot quantification of Trx80 protein levels in 3- and 10-month old wild-type and App^*NL-G-F*^ mice cortical homogenates (3-month old, n=5 wild-type, n=5 App^*NL-G-F*^; 10-month old, n=4 wild-type, n=5 App^*NL-G-F*^). Band intensities were normalized against tubulin. **E)** Western-blot quantification of Trx80 and Trx1 in 2-, 8- and 22-month old wild-type mice cortical homogenates (n=4 mice per group). Trx80 and Trx1 band intensities were normalized against actin. **F**) *Txn1* gene expression in neuron-specific bacTRAP-RNAseq data from different brain regions at 5-, 12- and 24-month old wild type mice. Brain regions include the layer II of the entorhinal cortex (EC2), hippocampal CA1, CA2 and CA3, somatosensory cortex (S1) and the dentate gyrus (DG). Statistical analysis between groups in the 2-, 8- and 22-month old wild-type mice brains was performed by one-way ANOVA for each protein. Statistical analysis between App^*NL-G-F*^ and wild-type mice was performed by Student t-test (*p<0.5; **p<0.01; ***p<0.001; **** p<0.0001).

To investigate the effect of amyloid pathology on Trx80 levels in the brain, we used APP^*NL-G-F*^ knock-in mice. This mouse model exhibits early amyloidosis and reactive gliosis at 2-3 months of age (21). We measured Trx80 protein levels in cortical samples from 3-month old (n=5 WT, n=5 APP^*NL-G-F*^) and 10-month old (n=4 WT, n=5 APP^*NL-G-F*^) female mice by western blot. Our results show that Trx80 is increased in 3-month old APP^*NL-G-F*^ mice compared to wild-type controls (p<0.0001, **Figure 1c**). However, Trx80 levels were decreased (p<0.05) compared to controls in 10-month old APP^*NL-G-F*^ mice (**Figure 1d**) resembling the decrease previously described in human AD brains (15).

To test whether aging, the most significant non-genetic risk factor for late-onset AD (22) affects Trx80 levels in the brain, we analyzed Trx1 and Trx80 protein levels in 2, 8 and 22 months old wild-type (WT) male mice. Western blot analysis of cortical homogenates did not show significant changes in Trx1 or Trx80 levels between 2 and 8 months of age. At 22 months, Trx1 levels decreased, while Trx80 levels increased compared to 2- and 8-month old mice (p>0,05, p<0.01 respectively, **Figure 1e**). While Trx80 increases 1.7-fold between 8- and 22-months of age, Trx1 decreases 0.5-fold, what suggests additional mechanisms, other than cleavage of the precursor, that specifically upregulate Trx80 at the protein level.

To further clarify *Txn1* expression levels in different brain regions and its changes during the mouse lifespan in a cell-specific manner, we revisited publicly available neuron-specific bacTRAP-RNAseq data from 5-, 12- and 24-month-old wild type mice (alz.princeton.edu) (23). As shown in **Figure 1f**, *Txn1* gene expression is higher in neurons of AD-relevant and memory-associated brain regions, such as principal neurons of the layer II of the entorhinal cortex (ECII neurons, the first neurons to degenerate in AD) and CA1, CA2 and CA3 pyramidal neurons in the hippocampal formation, compared to pyramidal neurons of the primary somatosensory cortex and granule neurons of the dentate gyrus. *Txn1* expression is significantly decreased in ECII neurons between 5 and 24 months of age (log FC = -0.47, FDR = 0.036) (23).

### Oxidative stress induces Trx80 production by neurons

Considering that Trx80 is the cleavage product of the antioxidant Trx1, we hypothesized that oxidative stress, a common event in aging and early amyloid pathology, might modulate Trx80 production. To test this hypothesis, we treated neuronal primary cultures with rotenone, a known inducer of oxidative damage (24-26). As expected, western blot analysis of neurons treated with 100 nM rotenone for 18h showed increased levels of nuclear factor erythroid 2-related factor 2 (NRF2), a well-known redox homeostasis regulator (27) (**Figure 2a**). Trx1 is a transcriptional target of Nrf2 (28), and its levels, together with Trx80 were increased in rotenone-treated neurons compared to control (p<0.0001, p<0.001 and p<0.05, respectively, **Figure 2a**).

**Figure 2.**
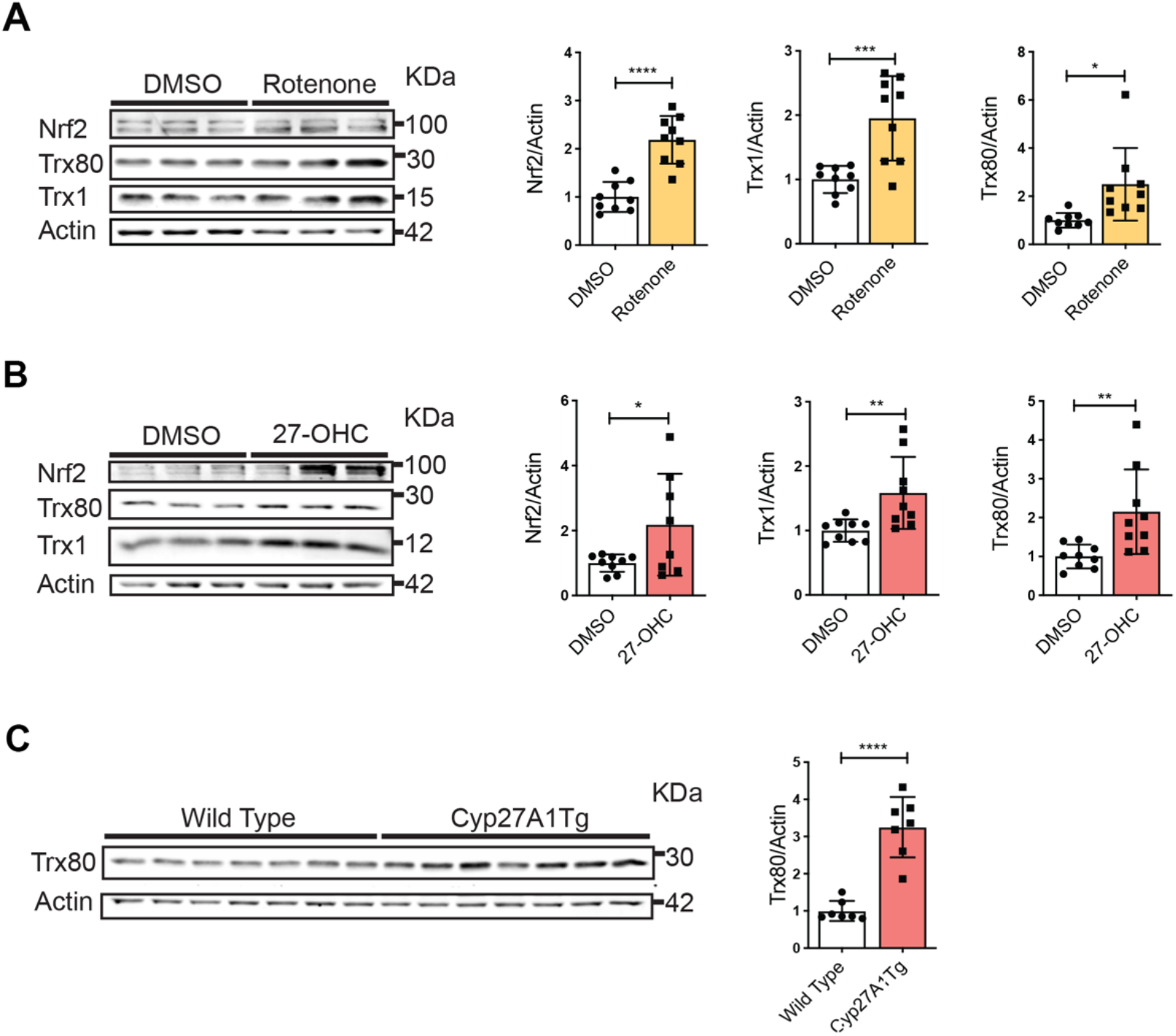
Rotenone and 27-hydroxycholesterol treatment induce an increased production of Trx80 in neurons. **A)** Western-blot analysis of neuronal primary cultures treated with 100 nM rotenone for 18h. Trx80, Trx1 and Nrf2 band intensities are normalized against Actin (n=3 biological replicates). **B)** Western-blot analysis of neuronal primary cultures treated with 1 μM 27-hydroxycholesterol for 2h. Trx80, Trx1 and Nrf2 are normalized against Actin (n=3 biological replicates). **C)** Western-blot quantification of Trx80 in 22-month old wild-type and Cyp27A1 overexpressing mice cortical homogenates (n=7 wild-type, n=7 Cyp27A1Tg). Results are shown as relative to controls. Statistical analysis was performed by Student t-test (*p<0.5; **p<0.01; ***p<0.001; **** p<0.0001).

To study this mechanism in more physiological conditions and *in vivo* we used the cholesterol metabolite 27-hydroxycholesterol (27-OHC). 27-OHC is a cholesterol metabolite that is elevated in AD brains (29) and causes oxidative stress (30, 31). We determined the effects of 27-OHC by profiling neurons 2h after 1μM 27-OHC treatment using RNA-sequencing. As expected, differential gene expression analysis (DGE) showed that 27-OHC modulates genes involved in cholesterol metabolism (**supplementary figure 1a** and **1b**) and pathway analysis revealed alterations in Nrf2-mediated oxidative response, among others (**supplementary figure 1c)**. Western blot analysis showed that Nrf2, Trx1 and Trx80 protein levels were increased in 27-OHC treated neurons compared to vehicle-treated controls (p<0.05, p<0.01 and p<0.01, respectively, **Figure 2b**). We next analyzed Trx80 levels in brain homogenates from a 27-OHC overproducing (Cyp27Tg) transgenic mouse model. These mice overexpresses the enzyme Cyp27A1, that converts cholesterol into 27-OHC, and hence have higher 27-OHC levels than wild-type controls (32) (33). Western blot analysis of Trx80 protein levels in cortical samples from 22 months old Cyp27Tg male mice (n=7) and age and sex-matched wild-type littermates (n=8) showed increased Trx80 levels in Cyp27Tg animals compared to controls (p<0.001, **Figure 2c**).

### Trx80 induces an interferon type-I response in microglia in a Trem2-dependent manner

To study whether Trx80 modulates microglial function, we treated mouse microglia in primary culture with 1μM recombinant Trx80 for 24h. RNA-sequencing followed by DGE analysis between Trx80 and vehicle-treated microglia revealed *Hpgd* as one of the most significantly downregulated genes in Trx80-treated microglia (**Figure 3a and 3b**). *Hpgd* codifies for 15-hydroxyprostaglandin dehydrogenase that participates in prostaglandins degradation and thus in the regulation of inflammatory response (34). Indeed, pathway analysis showed neuroinflammation and activation of interferon regulatory factors by cytosolic pattern recognition receptors as some of the top significantly upregulated pathways in Trx80-treated microglia, supporting the notion that Trx80 triggers a proinflammatory response in these cells (**Figure 3c**).

**Figure 3.**
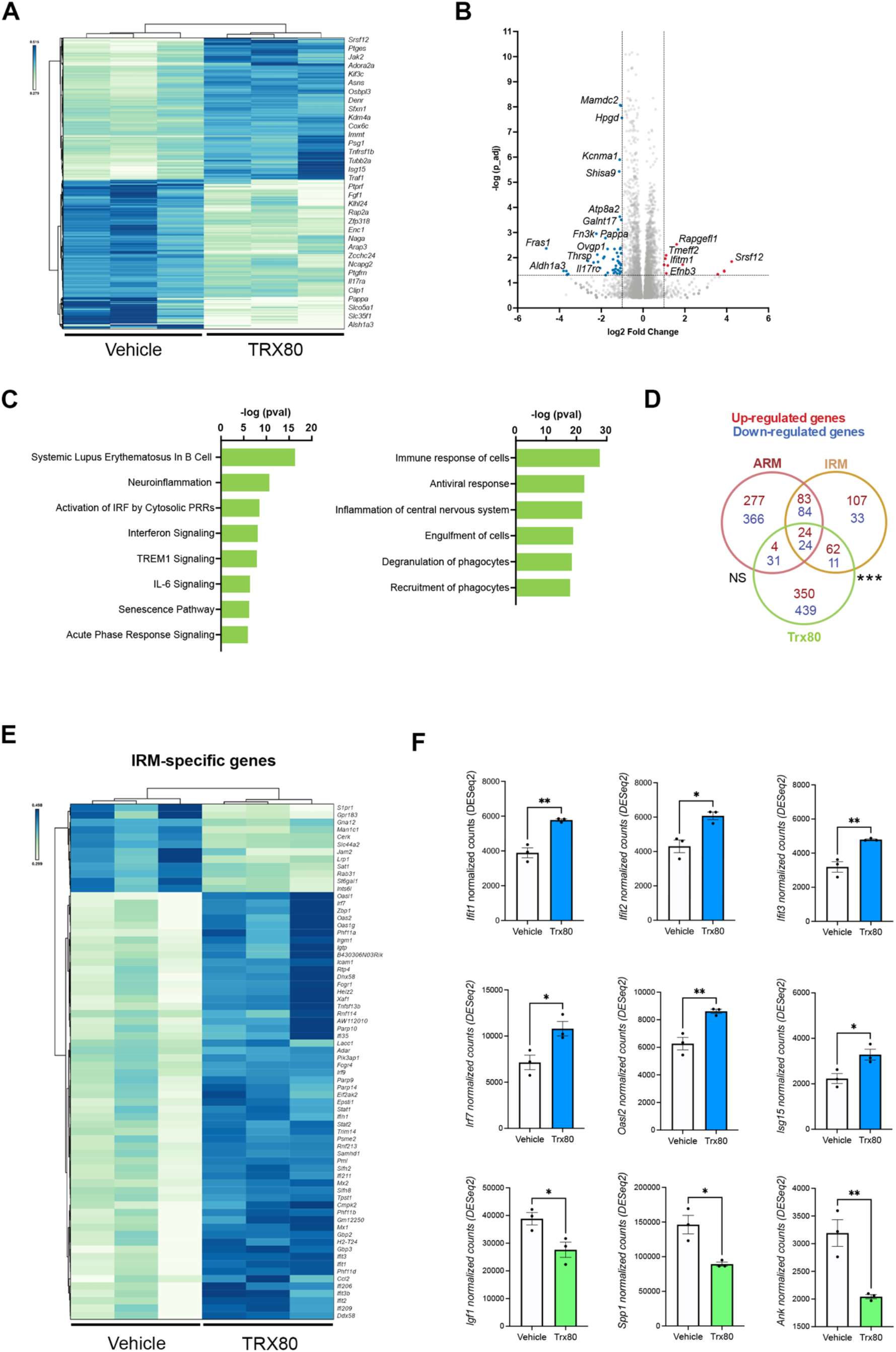
Trx80 induces an interferon type-I response in microglia. **A)** Heatmap of differentially expressed genes identified by RNA-seq (pval <0.05) between mouse primary microglia treated with 1 μM Trx80 for 24h and vehicle-treated controls. **B)** Volcano plot of the differential gene expression analysis between microglia treated with Trx80 vs vehicle (PBS) (cutoffs: padj < 0.05; log2 fold change > 1). **C)** Pathway analysis of Trx80-treated microglia versus vehicle. This analysis was performed with the Ingenuity Pathway Analysis (Qiagen) on genes that were differentially expressed (pval< 0.05). **D)** Venn diagram showing the overlap of downregulated (blue) and upregulated (red) DEGs in Trx80-activated microglia with the gene expression signatures (up and downregulated) of Activated Response Microglia (ARM, Red) and Interferon Response Microglia (IRM, Yellow). Statistical analysis was performed with a Fisher’s exact test (***p<0.0001). **E)** Heatmap of the expression levels of all the unique gene identifiers of IRMs in control and Trx80-treated microglia. **F)** Normalized counts (DESeq2) of *Ifit2, Ifit3, Irf7, Oasl2, Ifit1, Isg15, Igf1, Spp1* and *Ank* genes expression in Trx80-treated microglia versus vehicle-treated microglia (*p<0.5; **p<0.01).

To clarify whether Trx80 induces a reactive microglia phenotype, we compared the differentially expressed genes in Trx80-activated microglia with the gene signatures of ARMs and IRMs (9). The transcriptional profile of Trx80-activated microglia showed a highly significant overlap only with IRMs (**Figure 3d, 3e**). Specifically, we observed an increase in genes defined as IRM markers by previous studies (30), such as *Ifit2, Ifit3, Irf7* and *Oasl2*, as well as an increase in other interferon response-related genes (e.g. *Ifit1* and *Isg15*). *Igf1, Spp1* and *Ank*, described as upregulated markers of ARMs, where found to be decreased in Trx80-treated microglia compared to control (**Figure 3f**). *Ifit2, Ifit3, Irf7* and *Spp1* gene expression changes were confirmed in Trx80-treated microglia by RT-qPCR analysis in an independent experiment (p<0.05, p<0.001, p<0.0001 and p<0.001, respectively **supplementary Figure 2a**). This effect was not seen in Trx80-treated neurons (**supplementary Figure 2b**). Immunofluorescence staining of Ifit2, ISG15 and osteopontin (OPN, Spp1 protein) on Trx80 and vehicle-treated microglia showed higher Ifit2 and ISG15 protein levels (p<0.0001) and decreased Spp1 levels Trx80-activated microglia compared to control (p<0.001) (**Figure 4a, b and c**). Immunofluorescence analysis of ISG15 in 22 months old Cyp27Tg mice showed higher ISG15-positive microglia than age-matched wild-type controls (p<0.0001) (**Figure 4d**).

**Figure 4.**
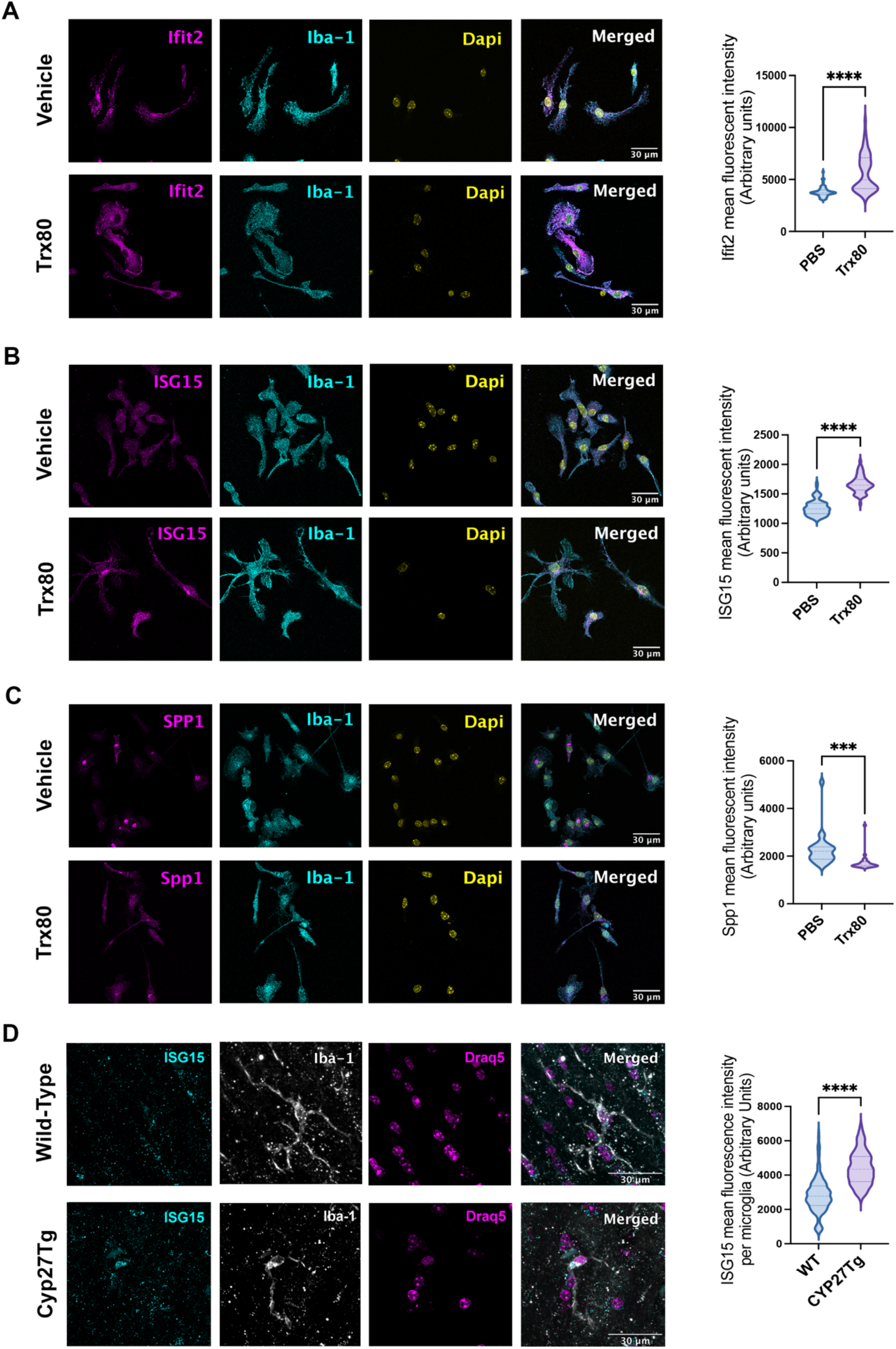
Increased Interferon related protein levels in Trx80-treated microglia. **A)** Immunofluorescence analysis and quantification of Ifit2 protein (magenta) in Trx80-treated (1 μM Trx80 for 24h) versus vehicle-treated microglia (cyan). Nuclei are stained with DAPI (yellow). **B)** Immunofluorescence analysis and quantification of ISG15 protein (magenta) in Trx80-treated microglia vs vehicle treated. **C)** Immunofluorescence analysis and quantification of Spp1 protein (magenta) in Trx80-treated microglia vs vehicle treated. Scale bar represents 30 microns. **D)** Immunofluorescence analysis and quantification of ISG15 protein (cyan) in 22 months old Cyp27Tg mice cortical microglia (grey). Nuclei are stained with Draq5 (magenta) Scale bar represents 30 microns. Statistical analysis was performed by Student t-test (***p<0.001),****p<0.0001).

We finally aimed to determine the molecular mechanism regulating Trx80-induced IRM phenotype. Trem2 is a transmembrane receptor that plays an essential role in microglia activation (3, 4) and its genetic variants have been associated with AD (5). Axl receptor tyrosine kinase is another transmembrane receptor that is upregulated in AD and described as a modulator of microglia-mediated phagocytosis of Aβ plaques (35) and pro-inflammatory signaling (36). To test whether the induction of an IRM phenotype in microglia by Trx80 is mediated by Trem2 or Axl, we knocked down each of these genes in mouse microglia in primary culture, followed by Trx80 treatment (**Figure 5a**). RT-qPCR analysis showed that partial Trem2 knock down prevents the Trx80-mediated increase in *Ifit2, Ifit3* and *Irf7* gene expression while it doesn’t affect *Spp1* gene downregulation. Knocking down Axl did not trigger significant differences in Trx80-induced expression of *Ifit2, Ifit3* and *Irf7* compared to control (scrambled shRNA transfected) microglia (**Figure 5b**). These results suggest that Trx80 effect on microglia interferon-response is Trem2-dependent.

**Figure 5.**
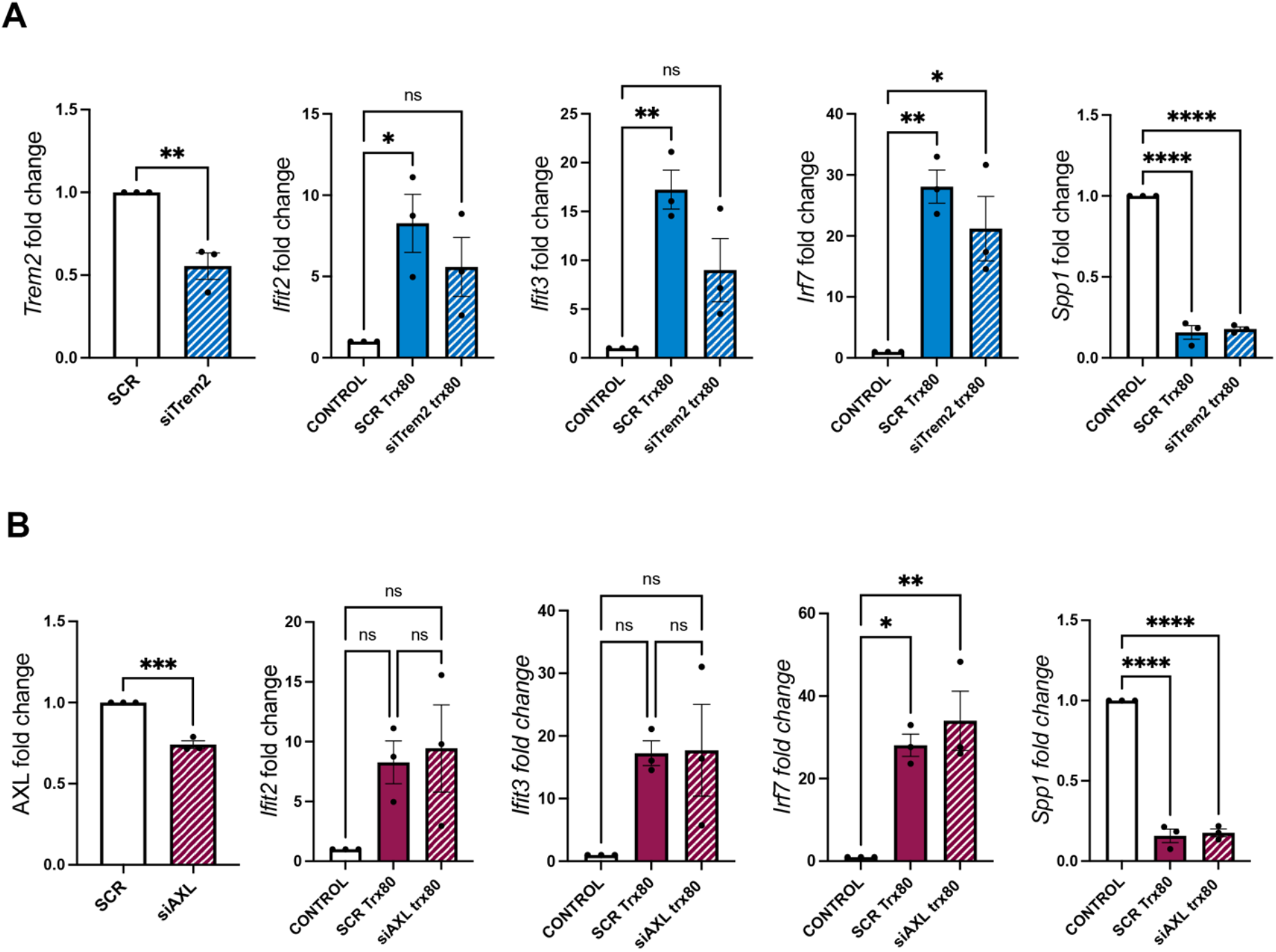
Trem2 silencing attenuates interferon-related gene up-regulation in Trx80-treated microglia. **A)** *Trem2, Ifit2, Ifit3, Irf7* and *Spp1* gene expression fold change in SCR Trx80 microglia (scrambled siRNA-transduced and Trx80-treated microglia) and siTrem2 Trx80 microglia (siTrem2 siRNA-transduced and Trx80-treated microglia) compared to their respective controls, that were given a value of 1 (scrambled siRNA-transduced, PBS-treated microglia and siTrem2 siRNA transduced, PBS treated microglia, respectively). **B)** *Axl, Ifit2, Ifit3, Irf7* and *Spp1* gene expression fold change in SCR Trx80 microglia (scrambled siRNA-transduced and Trx80-treated microglia) and siAxl Trx80 microglia (siAxl siRNA-transduced and Trx80-treated microglia) compared to their respective controls, that were given a value of 1 (scrambled siRNA-transduced, PBS-treated microglia and siAxl siRNA transduced, PBS treated microglia, respectively). Only one bar has been plotted as control for simplification purposes. Statistical analysis between groups was performed by one-way ANOVA (*p<0.5; **p<0.01; ****p<0.0001).

## Discussion

Trx80 peptide is found in the brain (15), where it prevents Aβ aggregation and inhibits Aβ toxic effects (15, 16). Despite its role on Aβ clearing, its biological function in the brain remains to be fully understood. Indeed, most of the studies on Trx80 have been performed in peripheral blood mononuclear cells, where it acts as a pro-inflammatory cytokine for monocytes, inducing their replication and enhancing the recruitment of lymphocytes (12-14). Innate immune pathways and neuroinflammation play an essential role in the pathogenesis of AD (37, 38). Therefore, in the present study we aimed to investigate the mechanisms regulating Trx80 synthesis in the brain, how they are affected by aging and AD-relevant pathological contexts and the effect of Trx80 on microglia function.

Trx1, Trx80 precursor, shows similar protein levels in all cell types: neurons, astrocytes and microglia. However, Trx1 mRNA levels were higher in neurons as well as Trx80 protein levels, pointing at neurons as the main source of Trx80 in the brain. These results suggest that neurons dedicate a part of Trx1 expression for Trx80 production under basal conditions. A previous study identified the metalloproteases ADAM10/17 as responsible for Trx1 cleavage into Trx80 (15). Our results suggest that the activity or the levels of these enzymes might be higher in neurons than in other cell types of the brain. This notion is supported by a previous study showing that ADAM10 levels are higher in neurons than in astrocytes (39).

We next analyzed Trx80 levels on APP^*NL-G-F*^ mouse brain homogenates, observing higher Trx80 levels at 3 months, followed by a decrease at 10 months of age. Interestingly, increased levels of Trx80 take place at the same age as the first Aβ depositions occur in this mouse model (21). This suggests that Trx80 production increases as an early stress response to Aβ accumulation, Aβ oligomers-induced alterations on neurons or secondary to inflammation induced by oligomers. Indeed, previous studies suggested that Trx80 is an early danger-response signaling molecule in the periphery (14). An explanation for its decrease at 10 months is that at this age, APP^*NL-G-F*^ mice show widespread Aβ depositions, prominent gliosis and synaptic loss, which could be translated into alterations in neuronal function, and possibly in the levels and/or activities of the metalloproteases responsible for the cleavage of its precursor (21, 40).

By analyzing cortical homogenates from wild-type mice at different ages, we showed that Trx80 levels increase with age, while Trx1 levels decrease over time. These results agree with previous studies of Trx1 and Trx80 levels on blood serum samples in a population of young and older adults (41). Trx1 levels decrease progressively throughout the mouse lifespan, while Trx80 increase after 8 months of age. These results support the notion that Trx80 levels are not only a consequence of the abundance of its precursor, but that they are regulated in a specific manner, most likely by the activity of the enzymes responsible for Trx1 cleavage (41) or its degradation.

Oxidative stress is a key factor in the cellular effects caused by aging (42-44). Elevated Aβ levels have been associated with oxidized byproducts of proteins, lipids, and nucleic acids in AD brains and *in vitro* Aβ treatments elevate intracellular ROS levels in neurons (45, 46). Therefore, we hypothesized that both aging and Aβ pathology could lead to increased Trx80 levels due to oxidative stress. Furthermore, alterations in the levels of cholesterol metabolites have been associated with AD (47) and 27-OHC, a cholesterol metabolite that is elevated in AD brains (29), leads to oxidative stress (30, 31). Here we show that both rotenone (*in vitro*) and 27-OHC (*in vitro* and *in vivo*) induce Nrf2 signaling and increased Trx80 levels in neurons. Interestingly, the cleavage of Trx1 into Trx80 renders the peptide devoid of its redox capacity (48). This, together with our results suggests that neurons increase Trx80 production under stress conditions with other purposes than mere antioxidant defense.

We next investigated the effects of Trx80 on microglia. Expression analysis on Trx80-treated microglia revealed that most upregulated pathways are associated with inflammatory response and processes related to viral and bacterial recognition and elimination. This transcriptional profile matches the effects of Trx80 reported in the periphery (14). Importantly, the transcriptional profile of Trx80-activated microglia shows significant similarities to the IRM subset of reactive microglia (9). The role of IRMs in the brain is still unknown. However, DNA damage promotes the production of type-I interferons (49) and IRMs have been found in the proximity to neurons harboring DNA damage (50), amyloid plaques containing nucleic acids (51), and to be enriched with age in the human brain (52). Altogether, these studies suggest that IRMs are associated with basal surveillance of neuronal stress and overall neuronal damage present in aging-related processes. Our results showing that Trx80 increases in the brain with age, suggest that Trx80 could be involved in the appearance of IRMs in aged brains. A microglia subset that is enriched in interferon response genes has recently been found in MCI and AD brains (8). Our results, showing an early increase in Trx80 in a mouse model of amyloid pathology, supports an association of this axis with AD pathology, however, whether the presence of interferon response microglia promotes or hinders the development of AD remains unknown. DAMs phenotype induction was previously shown to take place in two sequential steps: a Trem2-independent step, followed by a Trem2-dependent step (6). The mechanisms driving other phenotypes such as IRMs are still unknown. Here we provide evidence that Trx80 induces an IRM-like phenotype, and that this response is also dependent on Trem2. This finding suggests that IRMs and DAMs (9), although distinct and independent microglia phenotypes, are driven by Trem2 in combination with other signaling mechanisms that differ depending on the specific trigger and the biological context.

## Conclusions

Our findings identify Trx80 as a novel neuronal stress-induced signaling peptide that regulates microglia function. However, further work needs to be done to elucidate the potential contribution of Trx80 to the appearance of IRMs in different contexts and to fully understand the role of this microglia phenotype in health and disease.

## Abbreviations

27-OHC: 27-hydroxycholesterol;
Aβ: Amyloid-beta;
AD: Alzheimer’s Disease;
Ank: Ankyrin1;
ARM: Activated response microglia;
bacTRAP: bacterial artificial chromosome – translating ribosome affinity purification;
IRM: Interferon response microglia;
Isg15: Interferon-stimulated gene 15;
Ifit2: Interferon Induced Protein With Tetratricopeptide Repeats 2;
Irf7: Interferon Regulatory Factor 7;
Nrf2: Nuclear factor erythroid 2-related factor 2;
Oasl2: 2’-5’ oligoadenylate synthetase-like 2;
Spp1: Secreted Phosphoprotein 1;
Trem2: Triggering receptor expressed on myeloid cells 2;
Trx1: Thioredoxin-1
Trx80: Thioredoxin-80.

## Availability of data and materials

The datasets used and/or analyzed during the current study are available from the corresponding author on reasonable request.

## Authors’ contributions

J.G., P.R. and S.M. designed the study. J.G. acquired and collected the *in vitro* and *in vivo* data with the help of R.L.V., G.G. and M.L.. P.R., JP.R. analyzed the RNA-sequencing data. J.G. wrote the manuscript. A.C.M., P.R. and S.M revised the manuscript. All authors read, commented and approved the final manuscript.

## Acknowledgments

We would like to thank the Bioinformatics and Expression Analysis (BEA) core facility for their service and help with the RNA-sequencing analysis and Biomedicum Imaging Core (BIC) facility at Karolinska Institutet for their service with microscopy imaging.

## Ethics Approval

All procedures involving animals were approved by the local ethical committee in Stockholm (Ethical approval number: 4884/2019).

